# Rapid Detection of Sepsis using CESDA: the *Caenorabditis elegans* Sepsis Detection Assay

**DOI:** 10.1101/144873

**Authors:** Ling Fei Tee, Toh Leong Tan, Hui-min Neoh, Rahman Jamal

**Affiliations:** UKM Medical Molecular Biology Institute (UMBI), Universiti Kebangsaan Malaysia, Malaysia; Departmenf of Emergency Medicine, Faculty of Medicine, Universiti Kebangsaan Malaysia, Malaysia

**Keywords:** sepsis, *C. elegans*, chemotaxis assay, *C. elegans* Sepsis Detection Assay (CESDA)

## Abstract

Sepsis is a life-threatening condition which could be alleviated by rapid diagnosis and appropriate antibiotic administration. However, currently available laboratory tests for sepsis diagnosis lacks sensitivity and specificity; they also have long turn-around times. In this proof-of-concept study, the nematode *Caenorhabditis elegans* was used as a biological sensor to detect urine of sepsis patients in an assay designated as the *C. elegans* Sepsis Detection Assay (CESDA). From January to June 2016, 45 patients who were admitted to the Emergency Department of a university hospital due to suspected sepsis were included into the study. Urine samples were obtained from these patients and healthy controls and spotted onto CESDA assay plates. Subsequently, *C. elegans* were aliquoted onto the centre of the plates and allowed to migrate freely. Number of worms found in either spots or quadrants of the plates containing control or suspected sepsis samples were scored in 10 minute intervals in a 60-minute duration. The CESDA index was then calculated for each sample, where an index near +1 represented attraction of the worms towards the sample, while an index near -1 signified repulsion. Confirmatory diagnosis for suspected sepsis samples was determined using a combination of clinical criteria assessment and standard laboratory protocols. All patients who were positive for sepsis were found to have a CESDA index of > 0.1 (positive predictive value, PPV ≥87%). In addition, the worms were able to differentiate urine of sepsis patients from control as early as 20 minutes (p=0.012). Interestingly, the assay was also able to identify infection within 40 minutes of the test (AUROC = 0.80, p= 0.016). The rapidity of CESDA in sepsis and infection identification as well as the usability of urine samples which are non-invasive towards the patient in this method makes it an interesting protocol to be further explored for sepsis diagnosis.

## Introduction

Sepsis is a condition in which patients develop life-threatening single or multi-organ dysfunction due to dysregulated host response to infection [1]. Till today, the diagnosis of sepsis remains a challenge, as there is no single reliable test for its early confirmation or exclusion. Blood cultures offer low sensitivity, viral serology tests are costly, while the common laboratory screening parameters of white blood cell (WBC) count, erythrocyte sedimentation rate (ESR), and C-reactive protein (CRP) have poor sensitivity and specificity for diagnosis of sepsis [2-6]. Early sepsis diagnosis is important as it can help emergency medicine physicians perform risk stratification and initiate antibiotics promptly (if required), leading to better patient management and outcome [7].

*Caenorhabditis elegans* is a nematode widely used for studies in developmental biology and is a model organism for many diseases, particularly in neurobiology [8]. Recently, the nematode was reported to be able to sense and differentiate human cancer cell secretions, cancer tissues and urine from healthy control samples [9]. The chemotaxis assay, designated as the *C. elegans* nematode scent detection test (NSDT) works best on urine samples, with a sensitivity of 95.8% and specificity of 95.0%. Positive predictive value (PPV) and efficiency of the test were 67.6% and 95.0%, respectively.

It has been reported that cancer is the most common comorbidity associated with infection, and they share multiple similarities [10-12]. Inflammatory process mediated by T cells are expressed in both diseases. Persistent immune activation and inflammation leads to the activation of common signalling pathways that regulate immunity in both cancer and infection. In addition, inflammatory processes in both diseases lead to the release of similar pro-inflammatory and anti-inflammatory cytokines, increased levels of reactive oxygen and nitrogen species, tissue wasting, and increased apoptosis [12].

Although the olfactory molecule emitted from cancer urine samples that was sensed by the worms is still unknown, we suspect that the inflammatory process during an infection will also lead to emission of specific olfactory molecules that could be detected by *C. elegans* in urine samples of patients with infection, and perhaps, sepsis. To test this hypothesis, we proceeded to perform a *C. elegans* chemotaxis assay on urine obtained from sepsis patients and evaluated its sensitivity and specificity in detecting sepsis using urine samples obtained from patients admitted to Emergency Department, Universiti Kebangsaan Malaysia Medical Centre (ED-UKMMC). We designate this assay as the *C. elegans* Sepsis Detection Assay (CESDA).

## Materials and Methods

### Ethics statement

This study has been reviewed and approved by the Research Ethics Committee, Universiti Kebangsaan Malaysia with the ethics reference number: UKM PPI/111/8/JEP-2016-060.

### Study setting

This was a pilot, proof-of-concept study carried out in ED-UKMMC, from January 2016 until June 2016.

### Patient sampling and controls

The study population included patients aged 18 years or older who presented to ED-UKMMC during the study period with suspected sepsis based on the 2001 SCCM/ESICM/ACCP)/ATS/SIS International Sepsis Definitions Conference criteria [13]. Written informed consent was taken from all study participants; no minor was recruited into the study. Patients with suspected sepsis or infection were included into the study. Sepsis is defined as a condition where the patients has a minimum of two systemic inflammatory response syndrome (SIRS) criteria together with suspected infection. Patients with infection without fulfilling the SIRS criteria were classified as infection. Patients who were already on antibiotics, antiviral or antifungal drugs during their presentation to ED-UKMMC, patients who have cancer, were pregnant or with autoimmune diseases were excluded from the study. Confirmatory diagnosis for sepsis samples was determined using a combination of clinical criteria assessment and standard laboratory protocols. Control samples were obtained from healthy subjects who, at the time of study, were free from infection or cancer.

### Urine Sample collection

Approximately 5ml urine was collected from each study subject using a sterile urine container and stored at 4^o^C for not more than 48 hours until CESDA was performed.

### Worm cultures and bacterial strains

Wildtype N2 *C. elegans* strain was used for CESDA. For maintenance plates, the worms were cultured at 25°C under standard conditions on the Nematode Growth Medium (NGM) agar with *Escherichia coli* OP50 as their food source [14]. For assay plates, NGM agar was used without OP50 inoculation.

### C. elegans chemotaxis assay

The assay was conducted as described by Hirotsu et al with some modifications to the NSDT assay [9]. Briefly, urine samples were pre-warmed to room temperature prior spotting onto assay plates. Spotting design for the samples was slightly different from the NSDT assay, with the plate design as shown in Fig 1; test (sepsis, T) and control (c) samples were spotted onto different quadrants. In addition, spots where the urine samples were dispensed were also marked as points in the respective quadrants.

**Fig1.**
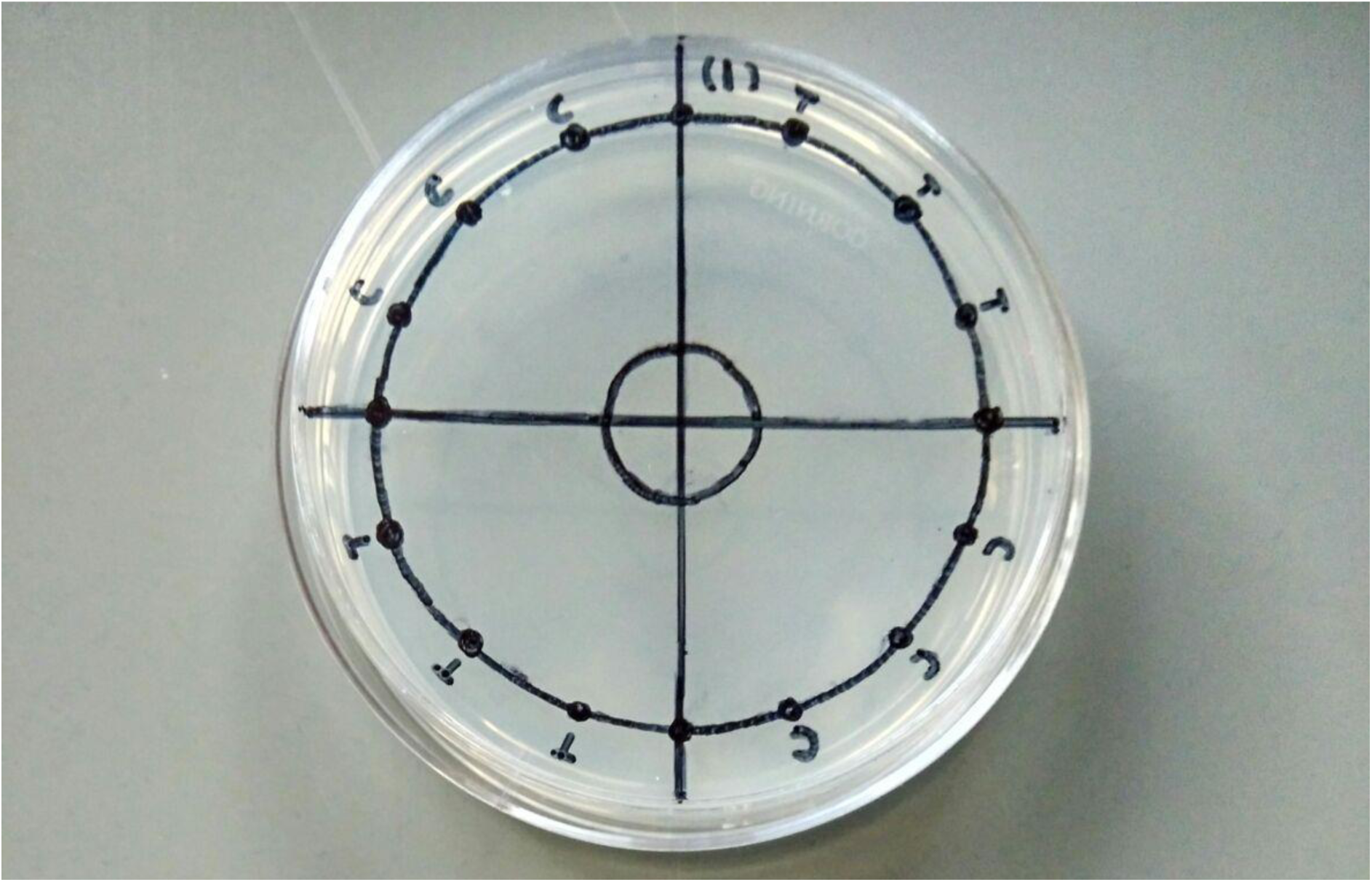
Plate design for CESDA. Urine samples were dispensed onto points (black-coloured full circles either labelled as “T” (sepsis) or “C” (control). Worms were then transferred onto a circle located in the middle of the assay plate and allowed to migrate. Number of worms (located in either “T” or “C” quadrants or at the exact points) were scored for each 10 minute-interval in a 60-minute assay.

Ten microliter of urine samples was spotted onto each point on the assay plate. Following that, 2ul of worms (about 50 worms) were transferred from a maintenance plate, washed with M9 buffer and aliquoted onto centre of the test plate. The worms were then allowed to migrate on the plate for 60 minutes. The number of worms found in the location of test and control quadrants as well as points were recorded for each 10 minute-interval for the 60 minute duration.

The CESDA index of the worms for each sample was then calculated as below:

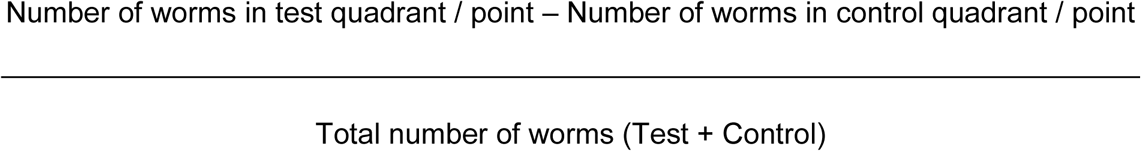

where a CESDA index near +1 represented attraction of the worms towards the sample, while a CESDA index near -1 signified repulsion [15].

### Statistical Analysis

Correlation of positive / negative chemotaxis index with sepsis / control samples was determined via Pearson’s chi-square test, where *p* < 0.05 was considered statistically significant.

## Results

During the duration of the study, a total of 166 patients were admitted to the ED-UKMMC for suspected infection. Out of this number, 56 subjects consented to the study and provided urine samples. Eleven patients were then excluded from the study: four had autoimmune diseases, two had anaphylaxis, two were already initiated antibiotics, one had malignancy and two were on long term steroid medication. The remaining 45 patients were eligible for this study based on the study’s inclusion and exclusion criteria. Among these, 36 patients’ urine samples were collected and tested in less than 24hours, while the remaining nine samples were collected and assayed between 24 – 48 hours. Ten patients had positive bacteria cultures. Table 1 shows demographic information of patients included into the study.

**Table 1.** Demographic data of patients recruited into the study.

|  | Total patients (n = 45) |
| --- | --- |
| Age (years; mean ± SD) | 45 ± 21 |
| Gender |  |
| Male | 31 (69%) |
| Female | 14 (31%) |
| Clinical Characteristic |  |
| Systolic Blood pressure | 130 ± 27 |
| Diastolic Blood Pressure | 77 ± 14 |
| Heart Rate (per minute) | 106 ± 16 |
| Respiratory Rate (per minute) | 21 ± 5 |
| Temperature ($^{\circ}$ Celsius) | 38.2 ± 1.0 |
| Total White Cell Count ($\times 10^9$) | 11.0 ± 9.1 |
| Sepsis | 34 (75.6%) |
| Infection | 39(86.7%) |
| Source of infection |  |
| Respiratory | 11 (24.4%) |
| Musculoskeletal | 6 (13.3%) |
| Urinary | 5 (11.1%) |
| Viral fever (non-specific) | 13 (28.9%) |
| Gastrointestinal | 1 (2.2%) |
| Blood/Catheter related | 2 (4.4%) |
| Positive bacterial culture | 10(22.2%) |

All patients who were diagnosed with sepsis were found to have a CESDA index of > 0.1 at 20 min (PPV = 87% for samples collected between 24-48 hours, and PPV = 92% for samples collected in less than 24 hours). Patients who were in the group of infection had a CESDA index of >0.232 at 40min (PPV= 92% for samples collected between 24-48 hours, and PPV=95.8% for samples collected in less than 24 hours) (Table 2). On further analysis, we found that CESDA could differentiate urine of sepsis patients from control as early as 20 minutes (p=0.012). The accuracy of the CESDA index was better for samples which were collected and tested in 24 hours compared to those between 24 to 48 hours (Table 3). We also found that the CESDA index which was calculated using worm numbers found on points rather than quadrants was associated more strongly with sepsis and infection (Tables 2 and 3). The ability of CESDA to predict sepsis and infection (for assays based on points) are shown in Table 4.

**Table 2.** CESDA index and AUROC values for urine samples collected and assayed in less than 48hours (n=45).

|  | **CESDA Index and AUROC** |  |  |  |
| --- | --- | --- | --- | --- |
|  | **Sepsis** |  | **Infection** |  |
| **Worm location,**<br>**time** | **Median ± SD** | **AUROC<sup>a</sup>**<br>**(95%CI)** | **Median ± SD** | **AUROC**<br>**(95%CI)** |
| Quadrant, 10min | 0.098±0.256 | 0.60 (0.44-0.77) | 0.095±0.254 | 0.66 (0.49-0.82) |
| Quadrant, 20min | 0.076±0.206 | 0.59 (0.41-0.78) | 0.053±0.221 | 0.62 (0.43-0.81) |
| Quadrant, 30min | 0.133±0.249 | 0.62 (0.47-0.78) | 0.123±0.251 | 0.59 (0.41-0.77) |
| Quadrant, 40min | 0.132±0.242 | 0.64 (0.43-0.84) | 0.123±0.239 | 0.55 (0.37-0.74) |
| Quadrant, 50min | 0.140±0.254 | 0.61 (0.41-0.81) | 0.137±0.278 | 0.60 (0.40-0.81) |
| Quadrant, 60min | 0.197±0.281 | 0.61 (0.43-0.79) | 0.194±0.304 | 0.58(0.38-0.77) |
| Point, 10min | 0.219±0.498 | 0.45 (0.26-0.64) | 0.224±0.494 | 0.45 (0.23-0.67) |
| Point, 20min | 0.452±0.390 | 0.67 (0.46-0.89) | 0.360±0.399 | 0.63 (0.32-0.94) |
| Point, 30min | 0.457±0.406 | 0.61 (0.41-0.81) | 0.458±0.394 | 0.70(0.47-0.93) |
| Point, 40min | 0.504±0.388 | 0.66 (0.47-0.86) | 0.508±0.371 | 0.80*(0.60-1.00) |
| Point, 50min | 0.567±0.405 | 0.56 (0.36-0.77) | 0.577±0.395 | 0.61 (0.33-0.88) |
| Point, 60min | 0.500±0.391 | 0.64 (0.46-0.83) | 0.364±0.565 | 0.82*(0.68-0.95) |

<sup>a</sup>AUROC = area under the receiver operating characteristic curve
\* indicates good accuracy (AUROC ≥ 0.80)

**Table 3.**
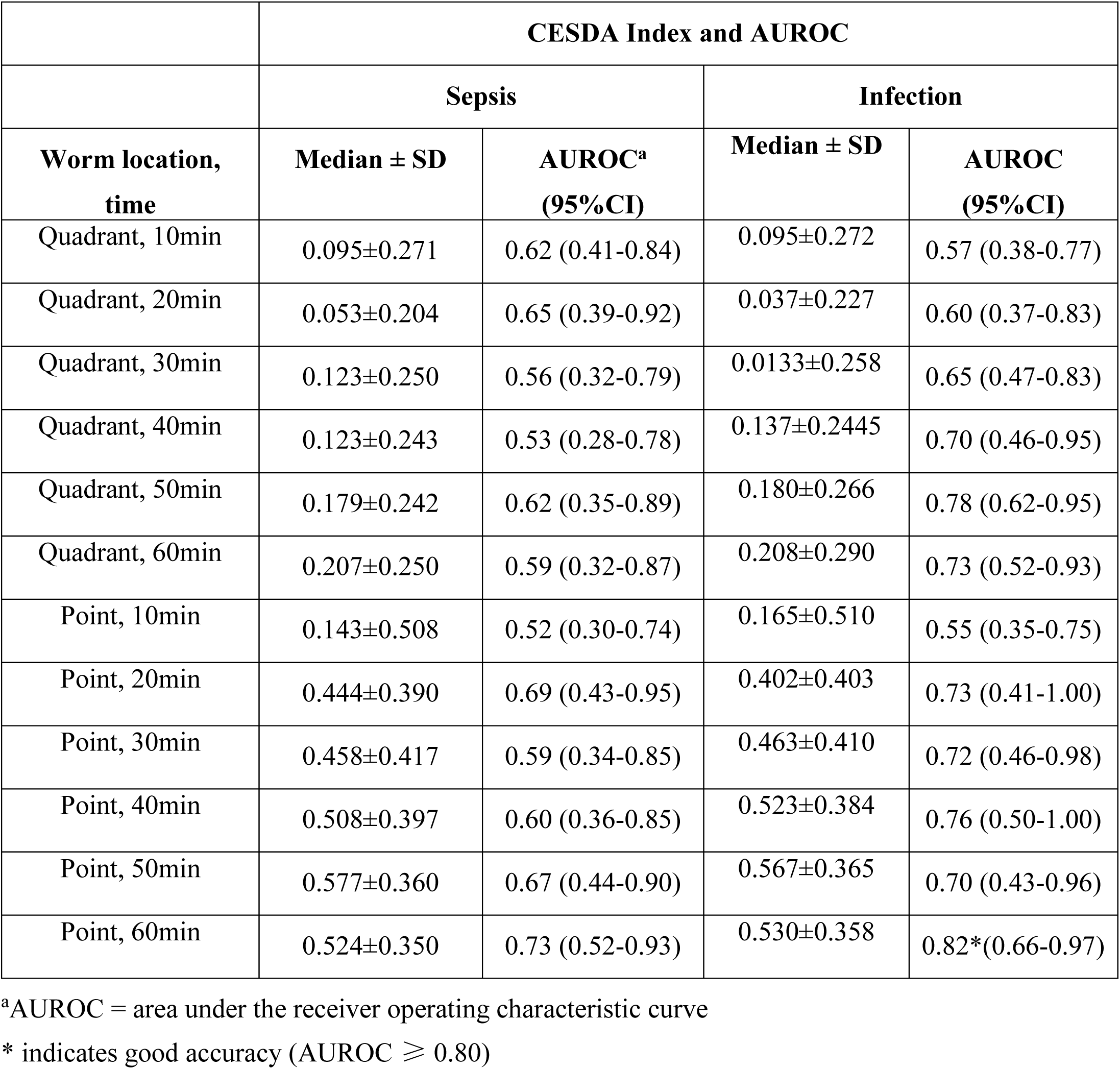
CESDA index and AUROC values for urine samples collected and assayed in less than 24hours (n=36).

|  | CESDA Index and AUROC |  |  |  |
| --- | --- | --- | --- | --- |
|  | Sepsis |  | Infection |  |
| Worm location,<br>time | Median ± SD | AUROC<sup>a</sup><br>(95%CI) | Median ± SD | AUROC<br>(95%CI) |
| Quadrant, 10min | 0.095±0.271 | 0.62 (0.41-0.84) | 0.095±0.272 | 0.57 (0.38-0.77) |
| Quadrant, 20min | 0.053±0.204 | 0.65 (0.39-0.92) | 0.037±0.227 | 0.60 (0.37-0.83) |
| Quadrant, 30min | 0.123±0.250 | 0.56 (0.32-0.79) | 0.0133±0.258 | 0.65 (0.47-0.83) |
| Quadrant, 40min | 0.123±0.243 | 0.53 (0.28-0.78) | 0.137±0.2445 | 0.70 (0.46-0.95) |
| Quadrant, 50min | 0.179±0.242 | 0.62 (0.35-0.89) | 0.180±0.266 | 0.78 (0.62-0.95) |
| Quadrant, 60min | 0.207±0.250 | 0.59 (0.32-0.87) | 0.208±0.290 | 0.73 (0.52-0.93) |
| Point, 10min | 0.143±0.508 | 0.52 (0.30-0.74) | 0.165±0.510 | 0.55 (0.35-0.75) |
| Point, 20min | 0.444±0.390 | 0.69 (0.43-0.95) | 0.402±0.403 | 0.73 (0.41-1.00) |
| Point, 30min | 0.458±0.417 | 0.59 (0.34-0.85) | 0.463±0.410 | 0.72 (0.46-0.98) |
| Point, 40min | 0.508±0.397 | 0.60 (0.36-0.85) | 0.523±0.384 | 0.76 (0.50-1.00) |
| Point, 50min | 0.577±0.360 | 0.67 (0.44-0.90) | 0.567±0.365 | 0.70 (0.43-0.96) |
| Point, 60min | 0.524±0.350 | 0.73 (0.52-0.93) | 0.530±0.358 | 0.82*(0.66-0.97) |

<sup>a</sup>AUROC = area under the receiver operating characteristic curve
\* indicates good accuracy (AUROC ≥ 0.80)

**Table 4.** CESDA index for predicting infection and sepsis for worms found on points.

| **CESDA index**<br>**for worms found**<br>**at “point”**<br>**location** | **CESDA**<br>**index**<br>**Cut off** | **AUROC** | **Sn,%**<br>**(95%CI)** | **Sp ,%**<br>**(95%CI)** | **PPV,%**<br>**(95%CI)** | **NPV,%**<br>**(95%CI)** | **PLR** | **NLR** | **Acc** | **Kappa** | **p\***<br>**(Fisher**<br>**'s**<br>**Exact**<br>**Test)** |
| --- | --- | --- | --- | --- | --- | --- | --- | --- | --- | --- | --- |
| **Urine < 48hours (n=45)** |  |  |  |  |  |  |  |  |  |  |  |
| Sepsis at 20min | 0.100 | 0.67 | 79<br>(62-91) | 64<br>(31-89) | 87<br>(75-94) | 50<br>(31-69) | 2.2<br>(1.0-4.9) | 0.3<br>(0.2-0.7) | 75.6 | 0.39 | 0.012* |
| Infection at<br>40min | 0.232 | 0.80 | 85<br>(69-94) | 67<br>(22-96) | 94<br>(84-98) | 40<br>(21-63) | 2.5<br>(0.8-7.9) | 0.2<br>(0.1-0.6) | 82.2 | 0.40 | 0.016* |
| **Urine < 24hours (n=36)** |  |  |  |  |  |  |  |  |  |  |  |
| Sepsis at 20min | 0.100 | 0.69 | 79<br>(60-92) | 71<br>(29-96) | 92<br>(78-97) | 46<br>(26-66) | 2.8<br>(0.9-9.1) | 0.29<br>(0.1-0.7) | 77.8 | 0.42 | 0.018* |
| Infection at<br>40min | 0.350 | 0.76 | 72<br>(53-86) | 75<br>(19-99) | 95.8<br>(81-99) | 25<br>(13-42) | 2.9<br>(0.5-16.0) | 0.4<br>(0.2-0.8) | 72.2 | 0.53 | 0.098 |

\* $p \leq 0.05$ indicates significant association, AUROC = area under the receiver operating characteristic curve; Sn = Sensitivity; Sp = Specificity; PPV = positive predictive value; NPV = negative predictive value; PLR = positive likelihood ratio; NLR = negative likelihood ratio; Acc= Accuracy; CI = Confidence Interval.

## Discussion

In this study, we explored the versatility of a *C. elegans* chemotaxis assay, which we designate as CESDA, to diagnose infection in patients admitted to ED-UKMMC. Interestingly, CESDA showed ability to differentiate sepsis patients from healthy controls and also from patients who had SIRS but not infection, and the detection can be completed as early as 20 minutes.

Our chemotaxis assay was a modification of the NSDT experiment employed by Hirotsu et al. (2015). The NSDT method was published in 2015 and the key finding was that it was able to differentiate the urine of 24 cancer patients from those of 218 healthy subjects. Indeed, the authors reported that five healthy subjects who were earlier identified as “cancer” according to the NSDT were later diagnosed with cancer within 2 years’ time of the study. The NSDT was found to be robust for various types and staging of cancer prediction, and has been suggested to be used as a cancer screening test [9].

In our study, CESDA index calculated using worm counts at points (our study) rather than quadrants on the assay plate (as in the NSDT protocol) provided a stronger association of the index with infection and sepsis. Stronger emission of olfactory molecules from the samples and thus attracting gathering of the worms in these points rather than the migration action of the worms across the quadrants might have increased the accuracy of the index in these points to predict infection and sepsis urine samples. In addition, we scored the CESDA index of the worms in 10-minute intervals for 60 minutes, compared to the NSDT protocol of observation at 60 minutes, and found that observation at 20 minutes provided a fair association of the index with sepsis, and a good association for infection at 40 minutes. Attraction of the worms towards distinct molecules emitted by either cancer, infection only or sepsis urine samples might have cause the differences observed in worm migration duration; this observation might be useful for future experiments to validate the chemotaxis assay.

With the modifications performed in our study, CESDA has the potential to be developed into a simple assay which could screen for and predict infection and sepsis in a span of 20 minutes. The rapidity of this screening test could provide clinicians and especially emergency medicine physicians the results they need in a shorter duration compared to the current diagnosis protocol, where serological tests and bacteriological cultures require 1-4 hours and more than 24 hours, respectively [7]. In addition, the chemotaxis assay uses urine as a diagnostic tool – this approach is beneficial especially for diagnosis in sepsis patients, as they are usually in hypovolemic shock where blood phlebotomy is complicated.

At present analysis, we could not state for sure that those patients who were positive for the CESDA index in our study will not be diagnosed with cancer in the future; however, they were cancer-free when they were included into this study. As mentioned in Hirotsu et al’s paper, the olfactory molecule being sensed by the nematodes to detect cancer is still unknown [9]. As we did not perform olfactory neuron ablation on the nematodes used in this study, with results from this study, we cannot conclude if the nematodes used in our study were attracted to the infected urine samples via olfactory sensing. However, as cancer and infection shares many T-cell-mediated inflammatory processes, it would be tempting to suggest that the worms might be also using their olfactory senses to detect olfactory molecules released due to activation of certain inflammatory processes in infection and sepsis.

Due to the explorative nature of this study, more samples need to be tested to validate further the nematodes’ attraction towards urine samples from infected and sepsis patients. Involvement of the nematodes’ olfactory system for their chemotaxis towards infected urine samples needs to be confirmed, and if indeed, the associated olfactory molecule needs to be identified.

## Conclusion

In this proof-of–concept study using samples from ED-UKMMC, urine samples of sepsis patients were found to be chemo-attractive to *C. elegans*. The rapidity of this assay for detecting life-threatening sepsis makes it worthy of further investigation; its usability should be validated in a larger cohort of samples.

## Acknowledgement

The authors would like to thank Dr Christabel Kang Wan Li, Dr Fatin Amirah bt Azmi, Dr Nadirah Hanim bt Abdu Hadi, Dr Dian Nasriana Nasuruddin and nurses of the ED-UKMMC for their help in patient recruitment and funding procurement.

